# Leaky evidence accumulation accounts for perceptual confidence and subjective duration

**DOI:** 10.1101/2025.03.17.643717

**Authors:** Ramla Msheik, Emma Sirouet, Simon Kelly, Nathan Faivre, Michael Pereira

## Abstract

Perceptual consciousness is defined as the subjective experience associated with the processing of sensory cues from the environment. Subjective experience unfolds over time and is accompanied by a sense of confidence, yet the mechanisms underlying these two properties remain elusive. Here, we propose a computational mechanism that accounts not only for the onset of subjective experience but also its subjective duration and associated feeling of confidence. Our model assumes that a percept becomes conscious when an ongoing, leaky accumulation of sensory evidence surpasses a perceptual threshold and stops being conscious when it falls back below the threshold due to leakage. Crucially, this perceptual threshold can be lower than the decision threshold for reporting the percept. Moreover, our model derives confidence in accurately detecting sensory evidence from the maximum reached by the evidence accumulation process following stimulus onset. We conducted a preregistered study in which three distinct models of evidence accumulation were fitted to behavioral reports of detection, confidence, and subjective duration during a face detection task under temporal uncertainty. The leaky evidence accumulation model accounted for the observed behavioral data best and outperformed alternative models without leakage. In a follow-up experiment, we investigated the impact of leakage adaptation on detection and subjective duration. We found that changes in leakage induced by contextual variations of face duration influenced both detection rates and subjective duration, as predicted by our model. Altogether, our findings suggest that leaky evidence accumulation is a suitable candidate mechanism to explain qualitative aspects of subjective experience, including subjective duration and feelings of confidence.

## Introduction

Subjective experience associated with sensory processing arises from the dynamic interactions between an individual and their ever-changing environment (Block, 1995; Nagel, 1974). As subjective experience is intrinsically private, most researchers rely on subjective reports (Francken et al., 2022) to assess its presence or absence (e.g. for weak stimuli). However, these subjective reports of stimulus detection only provide a limited account of the quality of subjective experience, characterized by specific temporal dynamics (Eagleman, 2008), as well as a subjective sense of confidence (Baranski & Petrusic, 1994). To investigate the temporal dynamics and confidence associated with subjective experience beyond detection, one can rely on the fact that subjective reports of duration and confidence do not strictly follow the physical properties of sensory stimuli, such as their duration or intensity. Subjective duration increases with stimulus intensity (Brigner, 1986), and confidence increases when the variance of the stimulus representation increases (Rahnev et al., 2012). Here, harnessing subjective judgments of confidence and duration, we propose a computational model based on leaky evidence accumulation explaining the presence or absence of subjective experience, its duration, and the associated feeling of confidence.

In perceptual decision-making, evidence accumulation models assume that noisy sensory evidence is integrated over time toward a *decision threshold*, the crossing of which leads to a decision triggering a motor response (Ratcliff & McKoon, 2008; Smith & Ratcliff, 2004). The *drift rate* is a parameter corresponding to the rate at which sensory evidence is integrated, and the *non-decision time* accounts for encoding and motor processes, that is, the part of response times that is not explained by evidence accumulation. Evidence accumulation was shown to successfully account for the speed and accuracy of a decision in various perceptual tasks (Gold & Shadlen, 2007; Hanks et al., 2014) in a biologically plausible manner (Cook & Maunsell, 2002; Donner et al., 2009; Hanks et al., 2015; O’Connell et al., 2012; Pereira et al., 2021; Shadlen & Newsome, 2001). A growing body of research suggests that the role of evidence accumulation might extend beyond decision-making to explain key properties of subjective experience, assuming that thresholds can be set on evidence accumulation not only for triggering action but also for triggering conscious perception. Notably, evidence accumulation was assumed to correspond to the unconscious processing of sensory signals towards a threshold for conscious detection, so a stimulus is detected when this threshold is crossed (Dehaene, 2011). This claim is supported by studies showing that the neural correlates of evidence accumulation can be observed irrespective of a corresponding motor plan (Bennur & Gold, 2011; Kumano et al., 2016; Twomey et al., 2016) and even in the absence of overt reports (O’Connell et al., 2012; Pereira et al., 2021; Stockart et al., 2024). While it is now clear that evidence accumulation subserves stimulus detection, its role in other aspects of subjective experience remains to be elucidated. Recently, we proposed that perceived intensity, duration, and confidence may be defined based on the time course of a specific form of evidence accumulation (Msheik et al., 2022; Pereira et al., 2022) that includes a *leakage factor* (Usher & McClelland, 2001). Put simply, a leaky integrator implements a low-pass filter that integrates over only a limited recent timeframe, thus rising when the stimulus is presented (i.e., when the drift rate is non-null) but falling back when the stimulus is removed, with limited buildup when there is only noise. Therefore, leakage offers an optimal policy for detecting stimuli when their onsets are temporally unpredictable (Glaze et al., 2015; Ossmy et al., 2013).

In the present work, we tested two main pre-registered hypotheses regarding the link between leaky evidence accumulation and aspects of subjective experience beyond stimulus detection (https://doi.org/10.17605/OSF.IO/8BV7N). First, we hypothesized that the confidence in correctly detecting a visual stimulus is proportional to the maximum level reached by the leaky accumulator following stimulus onset. Second, we hypothesized that the duration for which a visual stimulus is perceived consciously corresponds to the time during which the leaky accumulator remains above a perceptual threshold that might be distinct from the detection threshold. To test these two hypotheses, we designed a face detection task in which healthy volunteers attempted to detect face stimuli varying in intensity and duration. After each detection report, participants were asked to report their confidence in correctly detecting the face or the duration for which they perceived it. In experiment 1, we implemented a leaky evidence accumulation model that reproduced detection, confidence, and subjective duration. This model outperformed two alternative models without leakage. In experiment 2, we generalized our findings to a larger population. Finally, in experiment 3, we explored a model prediction for which leakage can be adapted dynamically depending on block-wise variations in stimulus duration, with consequences in terms of detection rate and subjective duration.

## Results

We used a face detection task during which participants were instructed to press a key on the computer keyboard upon perceiving a face. The stimuli, consisting of four artificial face images (two female, two male), were embedded in a series of 20 frames of visual noise presented every 200 ms. Each face appeared pseudo-randomly within each trial, starting at any frame between the fourth and tenth, and was presented for either 600 ms or 1 s. Face intensities were calibrated at the beginning of the experiment using an adaptive one-up/one-down staircase procedure to reach a 50% detection rate corresponding to the liminal intensity (Levitt, 1971). In each trial, the face intensity was either constant (static condition) or variable (dynamic condition) while present. The static condition included four face intensities corresponding to 0.8, 1, 1.2, and 1.4 times the participant’s liminal intensity. Meanwhile, in the dynamic condition, face intensity increased from 1x to 1.4x or decreased from 1.4x to 1x of the liminal intensity. 20% of trials contained no face (i.e. catch trials), in which the correct behavior was to refrain from making a response during the trial (no button press was required). After performing the detection task, participants were asked at the end of each trial to rate their confidence in their response (in confidence blocks) or to reproduce the duration of the perceived face (in duration blocks). In confidence blocks, participants used a continuous scale ranging from 0% (sure incorrect) to 100% (sure correct) to rate their confidence in their detection response; for missed faces and correct rejections, participants reported their confidence in having indicated the absence of a face. In duration blocks, we asked them to reproduce the subjective duration of the detected face following its offset by pressing and holding the computer mouse button for an equivalent duration.

Experiment 1 involved 4 participants performing eight experimental sessions each. This longitudinal design allowed us to assess how detection, reaction times, confidence, and subjective durations were affected by face intensity and duration, and to reproduce behavioral effects based on leaky evidence accumulation. We generalized our behavioral findings in experiment 2 using a cross-sectional design involving 20 participants performing one session each (supplementary information **S1**).

### Leaky evidence integration accounts for detection and response times

In the static condition of experiment 1, detection rates increased with duration as well as with face intensity, and the intensity effect was modulated by face duration (face intensity x stimulus duration: β = 0.15; *p* = 0.001). In the dynamic condition, long faces were better detected than short ones (β = −0.40; *p* < 0.001), but we found no effect of the face ramping profile (β = −0.02; *p* = 0.7) nor an interaction between face ramping profile and face duration (β = 0.017; *p* = 0.7). Higher face intensity in the static condition led to faster responses, with a stronger effect for short faces than long ones (face intensity x face duration: β = −0.03; *p* < 0.001). In the dynamic condition, faces with ramping-down intensities were detected faster than faces with ramping-up intensities, and this effect was stronger for short faces (face intensity x face duration: β = −0.029; *p* < 0.001). These behavioral results from four participants performing eight experimental sessions each were qualitatively similar to what we observed in experiment 2 (see **Figure 2B, 2D**, and **S1**).

**Figure 1:**
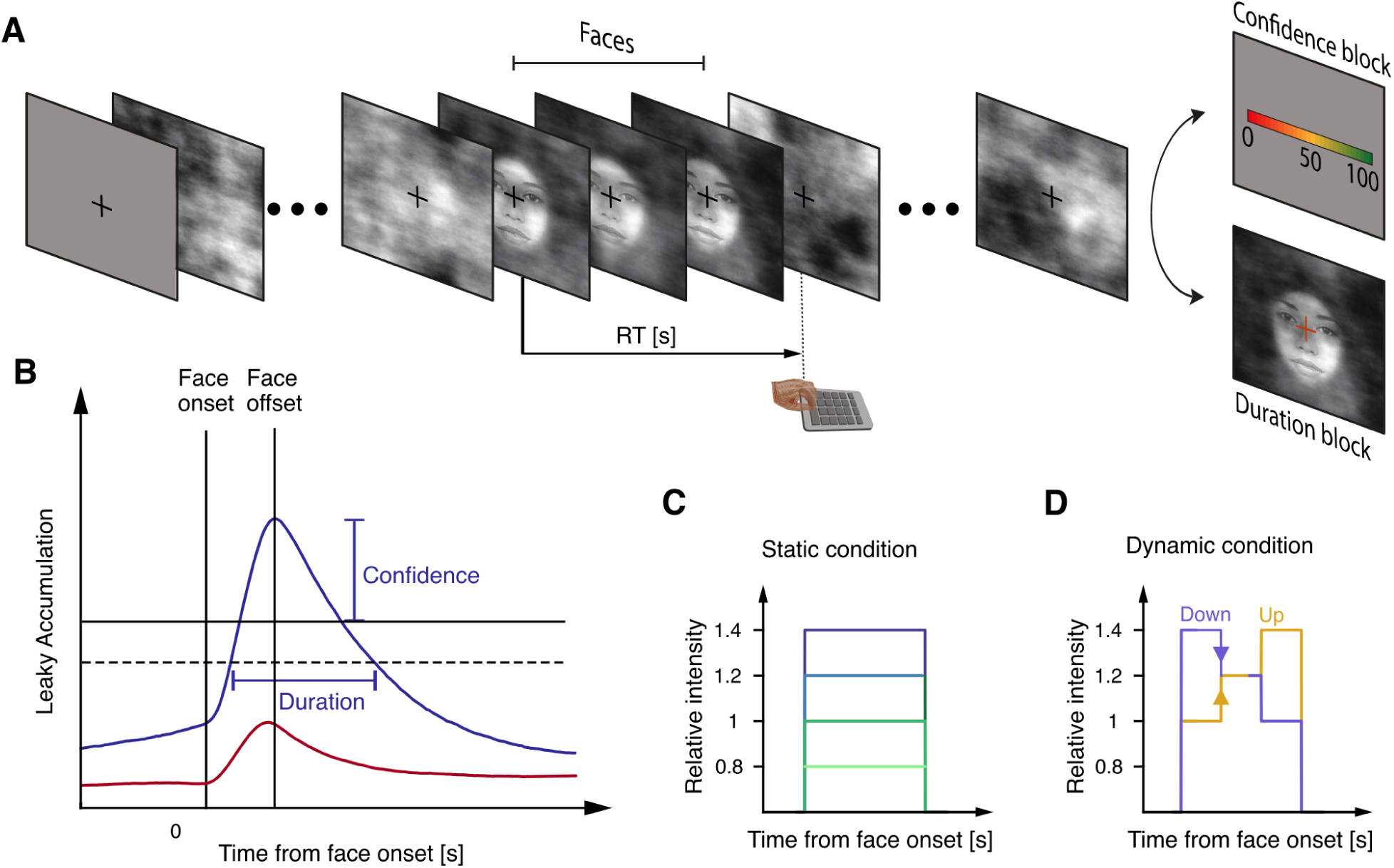
Experimental Paradigm and computational model. **A**. Experimental procedure. Participants were instructed to fixate on a cross and detect faces embedded in noise by pressing a key immediately upon detection. At the end of each trial, they were asked to assess their confidence using a continuous scale (confidence blocks) or to reproduce the duration of the detected face (duration blocks) by pressing and holding the computer mouse button for the same duration that they had previously perceived the face. As visual feedback a face was presented for as long as they held the button down. **B**. Example time courses of a leaky evidence accumulation process following the face onset, for a strong (blue) and weak (red) evidence trial. The leaky accumulator rises for as long as the face is present and then peaks and decays once the face offsets, and a ‘hit’ occurs when the process exceeds a decision threshold (solid), while a miss occurs when the threshold is not exceeded during the trial. Confidence is taken to scale with the maximum reached by the accumulation process. Meanwhile, subjective duration is defined as the time during which the accumulated evidence remains above a separate, lower, perceptual threshold (dashed line). Note: Non-decision delays are being ignored here for clarity of illustration. **C**. and **D**. Face intensity conditions. **C.** Static condition where face intensity remains constant throughout the trial, relative to each participant’s 50%-detection intensity when relative intensity = 1. **D**. Dynamic condition where face intensity increases from 1 to 1.4 times the 50%-detection intensity (yellow) or decreases from 1.4 to 1 times the 50%-detection intensity (purple).

**Figure 2:**
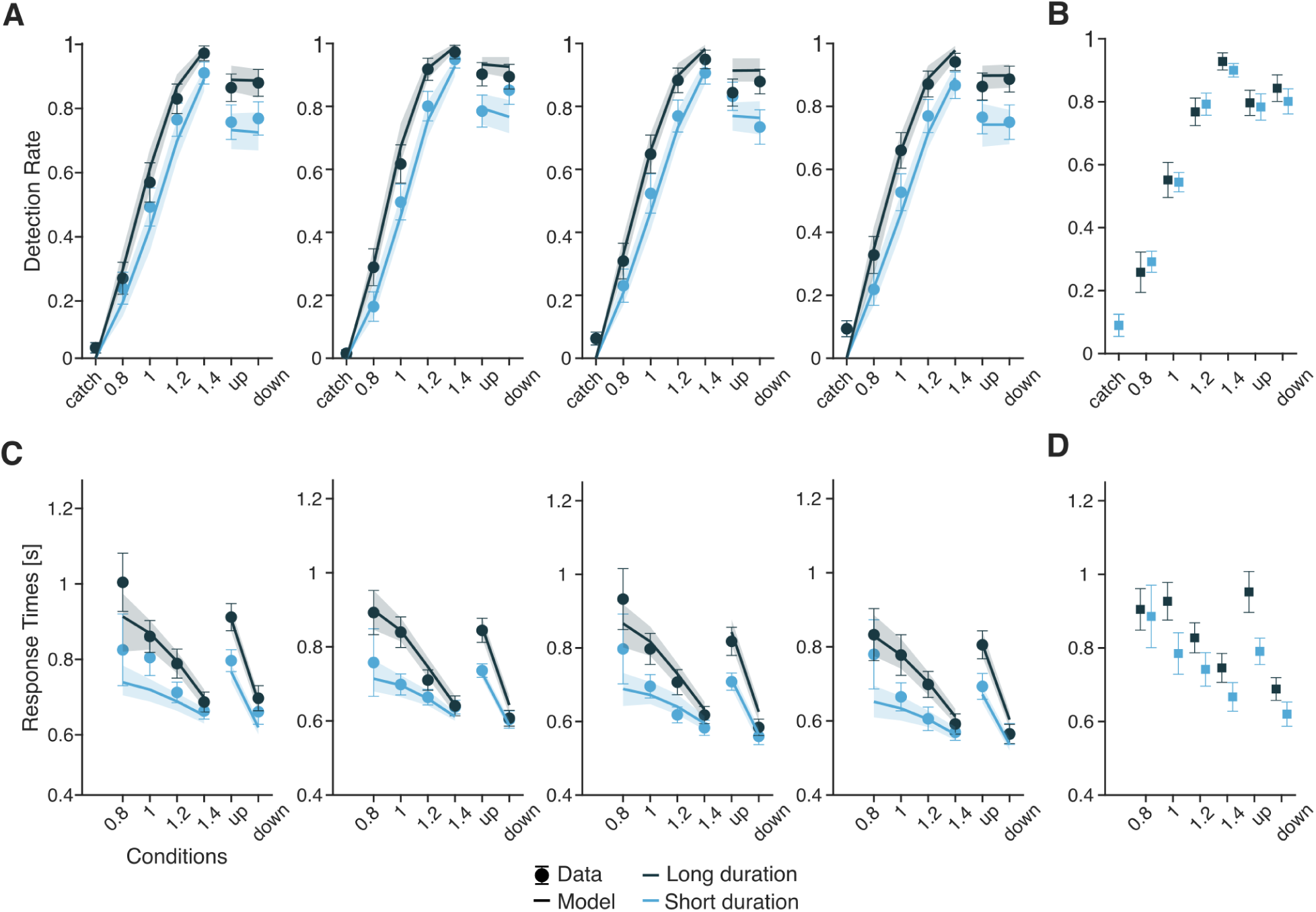
Detection results in experiments 1 and 2 with leaky evidence accumulation model fits. **A**. Individual detection rates for each of the four participants in experiment 1 (light and deep blue dots for short and long faces, respectively). Solid lines represent model fits for short faces (light blue traces) and long faces (deep blue traces), with shaded areas indicating 95% confidence intervals. **B**. Average detection rates across 20 participants for long faces (deep blue squares) and short faces (light blue squares) as a function of stimulus conditions in experiment 2. **C**. Individual response times in experiment 1. Solid lines represent model fits for short faces (light blue traces) and long faces (deep blue traces), with shaded areas indicating 95% confidence intervals. **D**. Averaged response times in experiment 2 for long faces (deep blue line) and short faces (light blue line) as a function of stimulus conditions.

Using only four free parameters (decision threshold, drift rate, non-decision time, and leakage), we successfully reproduced these behavioral effects by fitting a leaky evidence accumulation model to individual detection rates and reaction times. Leaky evidence accumulation was implemented as an exponential smoothing filter, with the leakage factor corresponding to the forgetting of past information. The leakage therefore corresponds to the decay constant which is inversely proportional to the time constant representing the integration time. The optimal time constant in terms of sensitivity corresponds to the stimulus duration (Ossmy et al., 2013). Of note, beyond optimality for sensitivity, stronger leakage reduces the delay introduced by the filter (see **S2**). We found that the fitted leakage values corresponded to short integration windows (26.7 ms, 49.4 ms, 22.6 ms, and 26.3 ms). Such short windows are suboptimal to discriminate signal trials from catch trials, but lead to quicker responses (**Figure S3**).

We compared the model fits against two alternative models: a perfect integrator model with no leakage and an extrema model (Hyafil et al., 2009; Stine et al., 2020)placing a decision threshold on instantaneous sensory evidence (see **Methods**). Our model captured observed results from all four participants (**Figure 2A, 2C**) better or similarly to the alternative models (see **Figures S4, S5, and Table S6**), and parameters had comparable values across participants (**Table S7**). This model comparison supports our hypothesis that detection relies on the temporal integration of noisy sensory evidence but with a strong leakage component, closer to an extrema model than a perfect integrator.

### Leaky evidence integration explains confidence

We fixed the parameters of the detection model and set out to test whether the dynamics of the leaky evidence accumulation process could explain a key aspect of subjective experience, namely the feeling of confidence with which participants perceived the face. In the static condition, we found that participants’ confidence judgments were modulated by an interaction between face detection and intensity (β = −0.02; *p* < 0.001). When participants reported perceiving the face, their confidence increased with both face intensity (β = 0.03; *p* < 0.001) and face duration (β = 0.01; *p* = 0.022). Conversely, when they did not perceive the face, confidence decreased with increasing face intensity (β = −0.01; *p* = 0.005) and longer face durations (β = −0.03; *p* < 0.02) (see **Figure S8**). In the dynamic condition, participants were more confident when they reported perceiving a face compared to when they did not (β = −0.06; *p* < 0.001), but — against our pre-registered predictions — we found no effect of face ramping profile (β = −0.007; *p* = 0.065) or face duration (β = −0.003; *p* = 0.41) on confidence judgments.

Following our pre-registered plan, we examined whether confidence was based on a readout of the maximum level reached by the accumulator during the trial. To do so, we simulated evidence accumulation traces using the model parameters found to fit detection rates and response times in experiment 1. For each simulated hit trial, we recorded the maximum level attained, and fitted a bias parameter and a multiplicative parameter to account for over- or under-confidence and confidence scaling (**Table S9**). We then saturated the result to the [0, 1] interval using a sigmoid function. Confidence corresponding to our model predictions captured the trend of behavioral data well (**Figure 3A**), emphasizing the role of evidence accumulation in accounting for confidence. Beyond average confidence, the model also accounted for confidence distributions (**Figure S10**). Our model could also reproduce the negative correlation between confidence and response times (supplementary information **S11**). On average, the maximum level of accumulated evidence was reached early after the detection response (individual mean: 114 ms, 141 ms, 125 ms, and 119 ms) and showed higher variance than a readout at a fixed time point after the decision.

**Figure 3:**
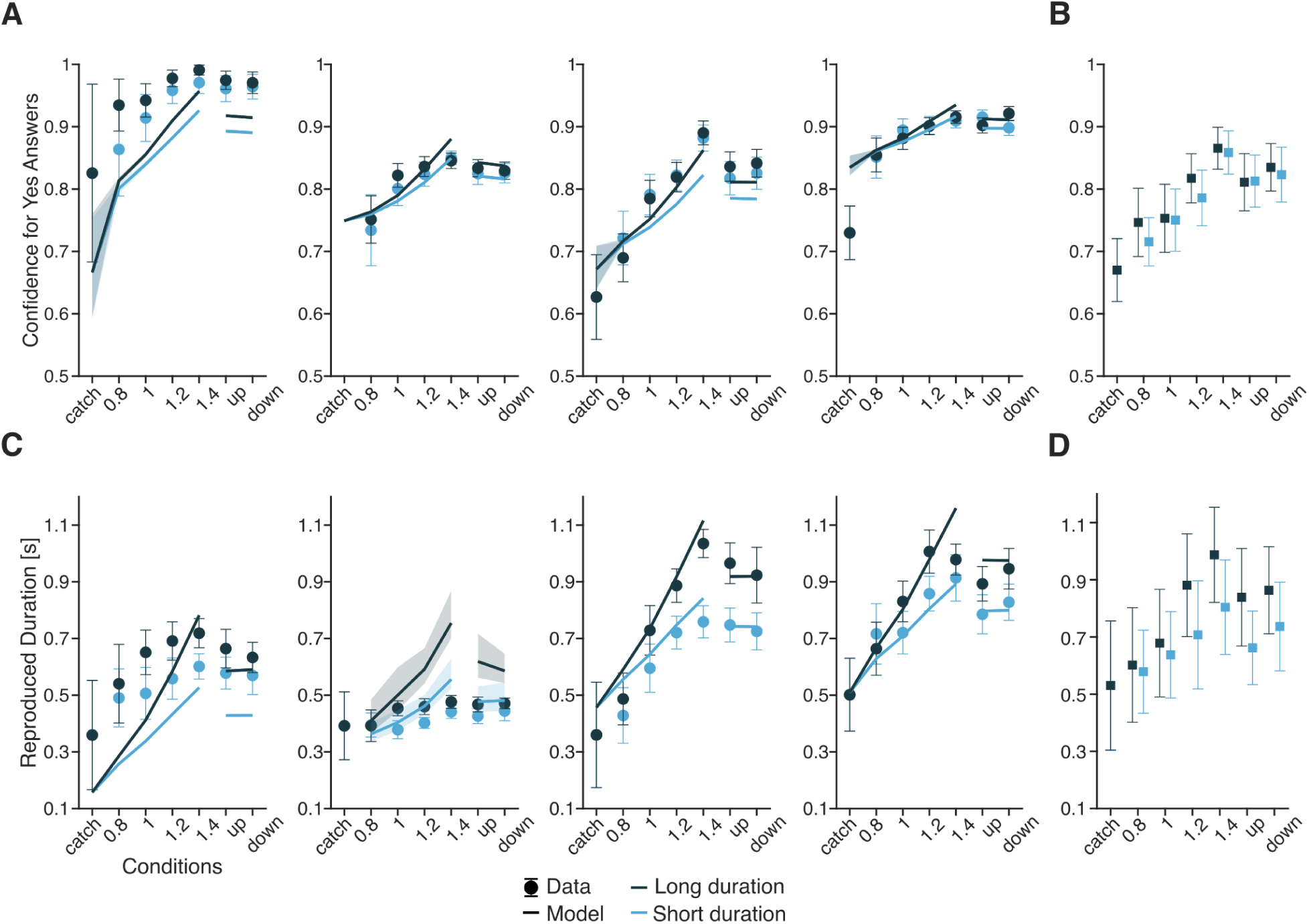
Confidence and subjective duration results in experiments 1 and 2. **A**. Individual confidence judgments for “yes” answers for four participants in experiment 1 (light and deep blue dots for short and long faces, respectively). Solid lines represent model fits for short faces (light blue traces) and long faces (deep blue traces), with shaded areas indicating 95% confidence intervals. Confidence data were scaled between 0 and 1 to allow comparison with the model’s predictions. **B**. Average confidence judgments for “yes” answers across 20 participants for long faces (deep blue squares) and short faces (light blue squares) as a function of stimulus conditions in experiment 2. **C**. Individual subjective durations for four participants in experiment 1. Solid lines represent model fits for short faces (light blue traces) and long faces (deep blue traces), with shaded areas indicating 95% confidence intervals. **D**. Average subjective durations across 20 participants for long faces (deep blue line) and short faces (light blue line) as a function of stimulus conditions in experiment 2.

One key advantage of defining confidence as the maximum level reached by accumulated evidence is the ability to model confidence without a decision (no decision threshold crossing in case of missed faces or correct rejections during catch trials). We applied the same readout of confidence for miss trials, thereby assuming that the closer the maximum accumulated evidence to the decision threshold during the whole trial, the lower the confidence in having been correct by reporting face absence. With this readout, we could reproduce the decrease in confidence in miss trials with increasing face intensity out-of-sample (**Figure S8**).

### Leaky evidence integration accounts for subjective duration

We then turned to subjective durations. In the static condition, participants reported longer durations as face intensity (β = 0.07; *p* < 0.001) and duration (β = 0.16; *p* < 0.001) increased, but we found no interaction between face intensity and face duration (β = 0.009; *p* = 0.5). In the dynamic condition, participants reported longer durations for long faces (β = −0.08; *p* < 0.001); however — against our pre-registered predictions — we found no effect of face ramping profile on the subjective duration (β = −0.001; *p* = 0.89).

We tested whether the dynamics of leaky evidence accumulation could relate to subjective duration. Based on previous findings, we preregistered that participants could have a conservative reporting bias for their detection reports and that their subjective experience of the face might relate to a *perceptual* threshold lower than the *decision* threshold used for reporting detection (Miller & Schwarz, 2006). Therefore, we modeled the perceptual threshold as a free parameter and estimated subjective duration as the time during which accumulated evidence stayed above this threshold plus an additive bias (**Table S12**). The pattern of predicted subjective durations reflected the observed behavioral trends of experiment 1, indicating that leaky evidence accumulation is a good candidate to explain subjective duration. Beyond average subjective duration, the model also accounted for subjective distributions (**Figure S13**). As for confidence, our model could also reproduce the correlation between subjective duration and response times (supplementary information **S11**).

### Adaptive detection behavior affects subjective duration

Having shown that leaky evidence accumulation is a plausible mechanism to explain subjective duration, we sought to test whether fluctuations in the environments leading to adaptations in stimulus detection would consistently affect subjective duration, as predicted by our model. First, we simulated detection rate responses and perceived durations based on the set of fitted parameters from participant P1 (Table **S7**). Then, we generated another set of detection rates and perceived durations using the same set of parameters but with a 4 % lower leakage value. The simulation results showed that higher leakage values led to lower detection rates and shorter perceived durations for both short and long stimuli, while lower leakage values produced the opposite pattern (**Figure 4A, 4B**). In a rapidly changing environment, fast detection is enabled by a short integration timescale, and this high leakage ensures a minimal rate of false alarms. Conversely, in an environment containing a majority of long-lasting face stimuli, it is more beneficial to integrate sensory evidence over longer timescales, with lower leakage (Ossmy et al., 2013). According to a model considering an overlapping mechanism for detection and subjective duration, this adaptation to longer timescales should lead to longer perceived durations.

**Figure 4:**
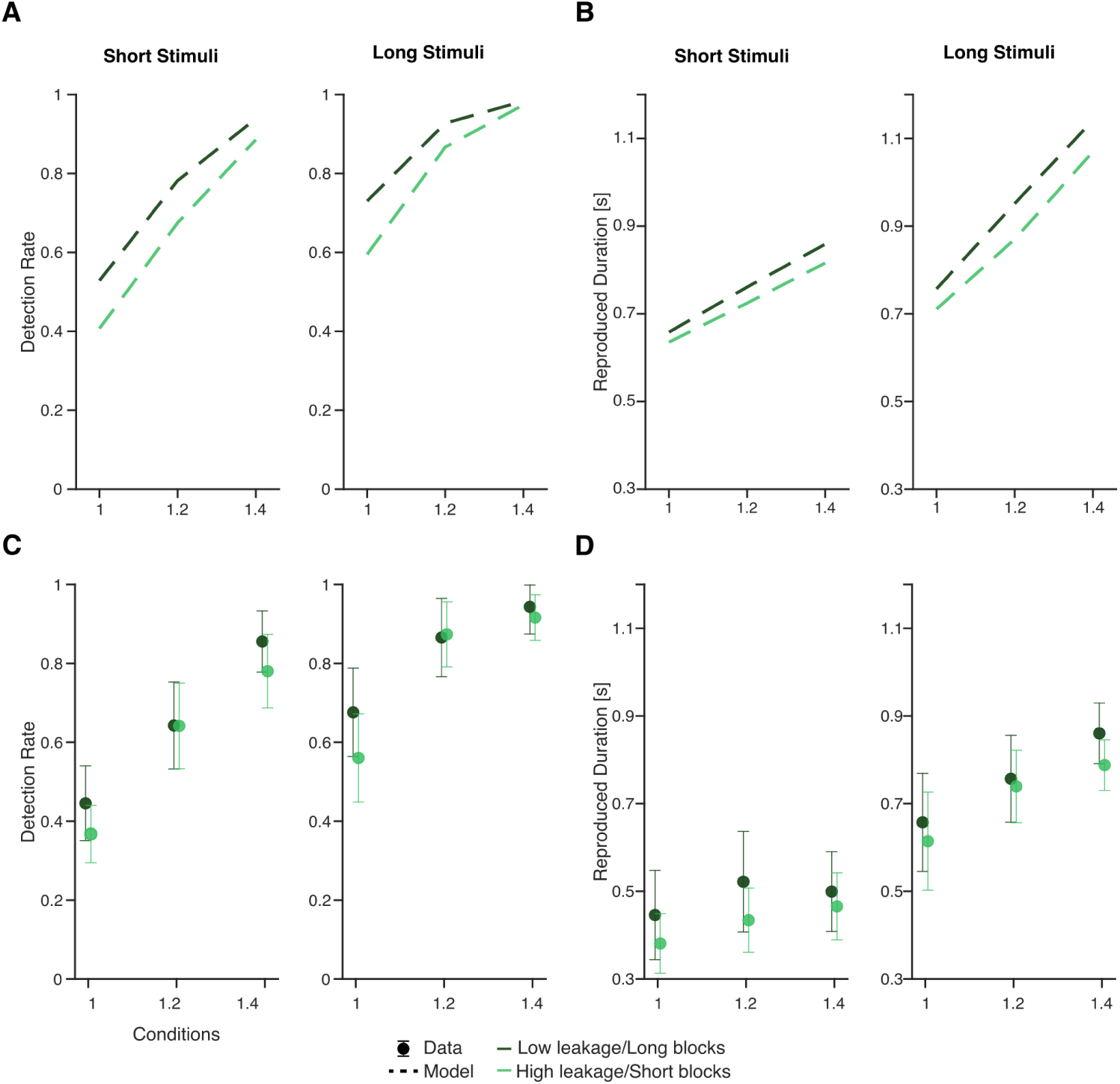
Model predictions and behavioral data for detection and subjective duration results in experiment 3. **A.** Model predictions for detection rates for short (left panel) and long stimuli (right panel) generated with high (light green dashed line) and low (deep green dashed line) leakage values. **B.** Model predictions for reproduced durations for short (left panel) and long (right panel) stimuli generated with high (light green dashed line) and low (deep green dashed line) leakage values. **C**. Average detection rate across 20 participants for short (left panel) and long (right panel) faces as a function of face intensity in predominantly short (light green dots) and predominantly long (deep green dots) blocks. **D**. Average subjective durations across 20 participants for short (left panel) and long (right panel) faces as a function of face intensity in predominantly short (light green dots) and predominantly long (deep green dots) blocks.

To test this hypothesis, we conducted a third experiment (experiment 3, N = 20), in which participants were required to detect and reproduce the duration of faces in blocks containing a majority of short or a majority of long faces. In short blocks, 75% of the faces lasted 600 ms, while 25% lasted 1 second. In long blocks, this proportion was reversed. As for previous experiments, detection rates increased with face intensity, and this effect was modulated by face duration (face intensity x face duration: β = −0.75; *p* = 0.008), with higher detection rates for long faces. Additionally, we found that participants detected faces better in predominantly long blocks compared to predominantly short ones (**Figure 4C**), both for short (β = 5.5; *p* < 0.001) and long faces (β = 8.5; *p* < 0.001). We compared the false alarm rates between short blocks (4%, s.d. = 0.017) and long blocks (4.83%, s.d. = 0.032). A Wilcoxon test showed no significant difference between the two types of blocks (p = 0.77). As for subjective duration, participants reported longer durations for high-intensity faces, and this effect was more pronounced for long than short faces (face intensity x face duration: β = −0.17; *p* = 0.002). Importantly, we verified our hypothesis as block type had a significant effect on subjective duration, with significantly longer subjective durations in long blocks compared to short ones for short faces (β = 0.09; *p* = 0.002) (**Figure 4D**). We note, however, that this effect was not found with a model comprising a random slope for the effect of block type, which corresponded to the best Bayesian Information Criterion (BIC).

## Discussion

We investigated whether a leaky evidence accumulation model for detection rates and response times could explain additional aspects of subjective experience, such as the feeling of confidence and subjective duration. To that end, we designed a face detection task in which participants reported the presence of a face and either rated their confidence in having accurately detected the face (confidence blocks) or reproduced the duration of the perceived face (duration blocks). We found that a model trained to reproduce detection rates and response times could parsimoniously account for subjective duration and confidence, suggesting that thresholded, leaky evidence accumulation plays an important role not only in gating access to consciousness (Cohen et al., 2020; Dehaene et al., 2014; Kang et al., 2017; Moutard et al., 2015; Pereira et al., 2021; Stockart et al., 2024) but possibly also in determining how subjective experience unfolds in time (Pereira et al., 2022). We first discuss the decisional part of the model that explains detection and response times, then turn to the subjective aspects the model can predict.

### Leakage: a key parameter for stimulus detection

Initially proposed to account for asymptotic constraints on the accumulation process (Usher & McClelland, 2001), leakage is formally defined as the forgetting factor in the accumulation process. In discrimination tasks, leakage has received relatively low empirical support compared to *bounded* evidence accumulation (Kiani et al., 2008). In detection tasks, however, leakage prevents noise accumulation in the absence of a stimulus, which would otherwise lead to false alarms (Ossmy et al., 2013) and the accumulation of irrelevant and outdated information, which could inflate the detection rate (see **Figure S4**). Here, we verified that leaky evidence accumulation better explained our data than perfect accumulation, confirming that participants discount past evidence. We also tested an *extrema* model for which the decision threshold is applied directly to noisy sensory information (i.e. with no evidence accumulation) and found that it provided either worse or similar fits as the leakage model (see **Figure S5**, **Table S6**). The optimal leakage factor to detect weak stimuli corresponds to an integration time constant that matches the physical stimulus duration. However, the leakage fitted by our model corresponded to a much shorter time constant, similar to previous studies using detection tasks (Ossmy et al., 2013). This contrasts with the long integration timescales found in the discrimination tasks typically used in the decision-making literature (Kiani et al., 2008). Even if suboptimal to discriminate or detect stimuli with predictable onsets, high values of leakage allow for shorter reaction times when stimuli are temporally unpredictable. The concept of leaky integration is consistent with neurophysiological data that supports a hierarchy of timescales across the cerebral cortex (Brincat et al., 2018; Chaudhuri et al., 2015; de Lafuente & Romo, 2006; Siegel et al., 2015), indicating that different cortical regions process information over different time windows, with shorter timescales (higher leakage) driving short-time (sensory) responses and longer timescales (lower leakage) informing strategic adjustments (Purcell & Kiani, 2016).

### Beyond stimulus detection: subjective confidence

We reasoned that leaky integration could explain subjective aspects associated with stimulus detection, such as confidence or subjective duration. Our results indicate that leaky evidence accumulation accounts for stimulus detection and explains the confidence in accurately detecting it. The model that best fitted our data allowed evidence to accumulate after crossing the decision threshold and linked confidence with the maximum reached by the accumulator. Our model differs from early works assuming a probabilistic readout of the state of the accumulators when the decision threshold is reached (Kiani et al., 2014; Kiani & Shadlen, 2009). This readout is only possible in discrimination tasks with an accumulator per choice. In detection tasks such as ours, there is only one accumulator whose state is constant when it reaches the decision threshold. There is strong evidence that some type of *unbounded* (also referred to as *post-decisional*) evidence accumulation occurs (Desender, Ridderinkhof, et al., 2021; Murphy et al., 2015; Resulaj et al., 2009) and that it might be used to account for confidence (Desender, Donner, et al., 2021; Goueytes et al., 2024; Pereira et al., 2020; Pleskac & Busemeyer, 2010). Post-decisional models of confidence have assumed that confidence is readout after a fixed amount of time (Desender, Donner, et al., 2021; Pereira et al., 2020; Pleskac & Busemeyer, 2010) or when a high confidence threshold is reached (Denmat et al., 2024; Grogan et al., 2023; Herregods et al., 2023). Our maximal evidence readout is compatible with the latter option but applied to a continuous confidence scale. The strong leakage factor combined with conservative decision thresholds (leading to a low false alarm rate) ensure that the readout happens soon after the decision threshold is reached (< 200 ms), similar to what was observed in electrophysiological studies (Desender, Donner, et al., 2021; Stockart et al., 2024). As maxima can only be defined a posteriori, this model of confidence also predicts that confidence evolves over time and can be revised (van den Berg et al., 2016).

Our results also confirmed that confidence co-varies with perceptual evidence and reaction times (Kiani et al., 2014; Rahnev et al., 2020). Specifically, we observed a negative correlation between confidence in detected stimuli and reaction times across all stimulus conditions. This result derives naturally from our postdecisional evidence accumulation model: if the decision threshold is crossed early, evidence can accumulate further above the decision threshold, leading to a high maximum evidence level, which results in high confidence ratings. Finally, reading out the maximum level of accumulated evidence allowed us to reproduce the decrease in confidence associated with increasing stimulus intensities when participants failed to detect a stimulus. This feature is impossible with a readout occurring at a particular time after the decision threshold is crossed, as no such event occurs for missed trials. Previous work from our group also showed that this confidence readout was better at describing confidence using both behavior and single-neuron activity in the parietal cortex of a human participant (Pereira et al., 2021). Altogether, our results regarding confidence highlight that detection and confidence formation can rely on the same mechanism of leaky evidence accumulation and that confidence corresponds to the maximal level of accumulated evidence.

### Beyond stimulus detection: subjective duration

Previous works have shown dissociations between a stimulus’ physical duration and the subjective duration of the corresponding percept. Most notably, stimulus intensity is known to impact subjective duration (Brigner, 1986; Eagleman, 2008), a phenomenon that we also observed in our data. As anticipated (Pereira et al., 2022), we could reproduce the subjective duration reported by participants based on the time between when accumulated evidence reaches a perceptual threshold and when it falls under it due to leakage. Although the leaky evidence accumulation model was trained only to reproduce detection behavior, it could reproduce the distributions of perceived duration with only two additional parameters to account for well known biases in reporting durations (Fraisse, 1984) and a possibly lower threshold for perception than for detection (Miller & Schwarz, 2006). In that sense, our model is parsimonious as it does not require additional assumptions, such as differences in the rate of an internal clock or a common magnitude system (Matthews & Meck, 2016). Moreover, it does not require dedicated accumulators for temporal coding (Ofir & Landau, 2022; Toso et al., 2021). Further model development will be needed to account for other temporal aspects of perception, including the capacity to retain stimulus in working memory or coding the subjective time between different events other than stimulus onset and offset.

Interestingly, our model did not only reproduce the average effects of stimulus intensity and duration on subjective duration. Indeed, both in the data and model simulations, response times explained a proportion of the variance of subjective duration that was not accounted for by the physical properties of the stimulus. This finding naturally results from the fact that response times are a proxy for the dynamics of the evidence accumulation process shared by both detection and subjective duration (as well as confidence). This result supports the view that stimulus detection and subjective duration share a common temporal integration mechanism subjected to distinct perceptual threshold and decision threshold. This claim was further supported by experiment 3, where changes in the stimulus environment, such as the proportion of long faces, affected both stimulus detection and subjective duration, although the physical properties of the faces were left unchanged.

### Subjective experience

Detected stimuli are associated with subjective experience, commonly referred to as “what it is like” to see them (Nagel, 1974). Building on findings from the decision-making literature, several theoretical accounts suggest that evidence accumulation may extend beyond decision formation and play a crucial role in the emergence of subjective experience (Dehaene, 2011; Pereira et al., 2022). Notably, the global neuronal workspace theory posits that a percept becomes conscious when the neural activity associated with it is made globally available to various cognitive processes such as reasoning, planning, and verbal reporting. This broadcasting occurs following an “ignition” process, which can be conceptualized as a threshold-crossing mechanism involving evidence accumulation (Dehaene et al., 2014; Moutard et al., 2015). Additionally, Shadlen and Kiani (2011) have suggested a close link between subjective experience and decision-making processes in the brain (Shadlen & Kiani, 2011). They propose that subjective experience might be mediated by a non-conscious “decision to engage” with the environment or a specific task. This view posits that mechanisms underlying decision formation share some fundamental features with those giving rise to subjective experience. It remained unclear, however, to what extent modeling detection reports could explain something about subjective experience beyond decision-making.

One behavioral study took advantage of the temporal variability of the onset of subjective experience and showed that an evidence accumulation model could reproduce the moment of the decision using a retrospective timing judgment (Kang et al., 2017). Here, we propose an integrated account of different phenomenological aspects associated with stimulus detection: subjective duration and associated confidence. With this approach, we aimed at mapping how perceptual experience unfolds over time, assuming that confidence judgment would relate to the strength of the perceptual experience and duration reproduction would relate to its duration. None of these behavioral measures are devoid of possible biases. Detection is believed to be conservative compared to subjective ratings of perceptual experience, and it is possible that participants did not report stimuli of which they had caught only a glimpse (Overgaard & Sandberg, 2021; Ramsøy & Overgaard, 2004). Response times contain a non-perceptual component that makes them a biased proxy to the onset of perceptual experience. Likewise, cognitive biases such as over- or under-confidence are expected to affect the confidence reported at the end of the trial, and it is unknown whether working memory is sufficient to correctly encode the strength and duration of the perceptual experience until it is reported. Nonetheless, by proposing an integrative account of these different behavioral proxies to perceptual experience and modeling generic biases for each of them, we believe that we will get closer to explaining the latent variable of perceptual experience.

Lastly, our approach poses constraints on the neural correlates of subjective experience or consciousness (Koch et al., 2016) as a true correlate of perceptual experience should not only distinguish perceived and non-perceived stimuli but should last for as long as perceptual experience lasts (Pereira et al., 2022). Some electrophysiological studies have started to consider the dynamics of the physical stimulus to further pinpoint the correlates of perceptual experience (Broday-Dvir et al., 2023; Hense et al., 2024; Podvalny et al., 2017; Vishne et al., 2023). In previous studies, we found neural activity with temporal profiles consistent with a leaky evidence accumulation process in the inferior temporal cortex during face detection (Stockart et al., 2024) and in neurons in the posterior parietal cortex for vibrotactile detection (Pereira et al., 2021). Future work will confirm whether such activity also covaries with subjective duration.

To conclude, we propose a parsimonious model to account for phenomenal aspects of subjective experience including perceptual confidence and subjective duration. While our behavioral and computational results support LEAP, further electrophysiological research is needed to elucidate how this mechanism is implemented in the brain.

## Materials and Methods

### Participants

Four participants (all right-handed, two female) participated in experiment 1, in which they completed eight sessions over 15 days (total: 14,336 trials). Twenty healthy individuals (aged 18-23, 18 right-handed, 16 female) with normal or corrected-to-normal vision participated in experiment 2. Two participants were excluded due to a high rate of false alarms (>30%; pre-registered). Twenty new healthy participants (aged 18-30, all right-handed, 15 female) with normal or corrected-to-normal vision participated in experiment 3. Two participants were excluded for exceeding the false alarm threshold (30%). The experimental protocol was approved by the local ethics committee of Grenoble Alpes University (Avis-2019-11-05-4) and pre-registered on the Open Science Framework platform (https://doi.org/10.17605/OSF.IO/8BV7N). All participants were recruited through an online advertisement on the laboratory’s website. They provided written informed consent and were financially compensated for their participation (10€/h).

### Apparatus and stimuli

Stimuli were displayed on a 15-inch Dell Precision 7550 laptop running at 60 Hz with a 1920 x 1080 pixels resolution. They consisted of four grayscale pictures (two male, two female) downloaded from https://generated.photos/ and cropped to contain exclusively facial features. They were presented at the center of the screen and represented 2.48° of the visual angle. Faces were embedded in noise patterns obtained by extracting the phase component of the Fourier transform from the face image, randomly permuting this phase, recombining it with the original magnitude component, and finally inversely transforming the result back into the spatial domain using the inverse Fourier transform.

In face-present trials, faces appeared at different intensities and lasted either three (600 ms) or five (1 second) frames with a different noise pattern for every frame. In experiments 1 and 2, face intensity varied randomly across trials in two different ways. In static trials, face intensity was constant and corresponded to 0, 0.8, 1, 1.2, or 1.4 times the individual liminal intensity. In dynamic trials, face intensity varied within the same trial, either increasing in a stepwise manner (from 1 to 1.4 times the liminal intensity, “ramp-up” condition) or decreasing (from 1.4 to 1 times the liminal intensity, “ramp-down” condition). In experiment 3, face intensity was constant and corresponded to 0, 1, 1.2, or 1.4 times the individual liminal intensity.

The individual liminal intensity was estimated through a one-up/one-down staircase procedure involving 50 trials (80 trials for experiment 3), using a relative intensity step of 5%. Then, the threshold value was calculated by averaging the intensities of the last five trials in the staircase. This adaptive procedure continued in experiments 1 and 2 to track the individual’s perceptual threshold throughout the rest of the experiment for short (600 ms) threshold stimuli (100%) corresponding to 1/13 trials.

### Procedure

All participants were seated at a 60 cm distance from the computer screen. The experiment was implemented using Matlab and the Psychtoolbox library (Brainard, 1997; Kleiner & al., 2007; Pelli, 1997). Following the staircase procedure, each session of experiments 1 and 2 included two confidence and two duration blocks, each consisting of 113 trials. Trials started with a fixation cross (10 pixels long and 2 pixels thick) presented on a gray background for 600 ms, persisting in subsequent frames. Afterward, a sequence of 15 stimulus frames was presented at 5 Hz for 3 seconds (i.e., each frame lasted 200 ms). In face-present trials, the face image appeared randomly (uniform distribution) after the fourth frame (600 ms post-sequence onset) and before the tenth frame (1800 ms post-sequence onset). Upon perceiving a face, participants were instructed to press the ‘2’ key on the numeric keyboard. A response was considered a hit if participants responded after the stimulus presentation and before the end of the trial. Otherwise, a failure to respond was considered a miss. The task following face detection varied between confidence and duration blocks.

In confidence blocks, participants were asked to assess their confidence in their responses using a continuous scale from 0% (sure incorrect) to 100% (sure correct). After the sequence of 15 frames, a continuous colored bar presented on a gray background appeared and participants indicated their confidence using the computer mouse.

In duration blocks, participants were asked to reproduce the subjective duration of the face they detected (i.e., no duration report was enforced following undetected faces). A screen of visual noise frames with a red cross at the center appeared, and participants were asked to reproduce the subjective duration of the detected face by pressing the computer mouse button as long as the duration for which they had perceived the face. As feedback, the face appeared at an intensity corresponding to 150% of the perceptual threshold as long as participants pressed the mouse’s button. The order of confidence and duration blocks was counterbalanced across sessions in experiment 1 and across participants in experiment 2.

In experiment 3, following the staircase, each session included three predominantly short and three predominantly long blocks, with each block containing 113 trials, 25% of which were catch trials (face-absent trials). In the predominantly short blocks, 75% of face-present trials featured short faces, and 25% featured long faces. Conversely, in the predominantly long blocks, 75% of the face-present trials featured long faces, and 25% featured short faces. Following stimulus detection, participants were requested to reproduce the perceived duration of the detected face. The order of predominantly short and predominantly long blocks was counterbalanced across participants, with participants starting either with the three predominantly short or three predominantly long blocks.

### Data analysis

We measured response times relative to face onset. Before analysis, responses classified as anticipatory (response times < 0 s; 0.91±0.67% in experiment 1, 2.24±2.62% in experiment 2, and 0.77±1.34% in experiment 3), early (response times < 0.2 s; 1.14±0.75% in experiment 1, 3.02±3.28% in experiment 2, and 0.98±1.69% in experiment 3), and late (response times > 2 s; 0.09±0.07% in experiment 1, 0.38±0.34% in experiment 2, and 1.02±1.49% in experiment 3) were excluded. Additionally, since we were interested in the confidence of being correct, we removed trials with confidence ratings below 50% (4.21±3.88% in experiment 1, 12.66±8.99% in experiment 2). While this excludes trials potentially involving changes of mind, these were not the focus of the present study.

In experiments 1 and 2, behavioral data were analyzed using mixed-effects (generalized) linear models with sum coding according to two main independent variables: the relative face intensity (continuous value between 0.8 and 1.4 in the static condition; *ramp-up* or *ramp-down* categorical variable in the dynamic condition) and the face duration (short or long). Data from static and dynamic trials were analyzed separately. A main factor’s effect was considered significant if the p-value was smaller than 0.05.

We used a generalized mixed-effect model (GLMM) with binomial distribution and a logit link function to evaluate the relationship between participants’ responses and face stimulus characteristics.

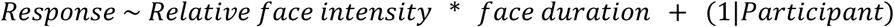

Response times and subjective durations were analyzed using generalized mixed-effect models (GLMM) with gamma distribution and logarithmic link function. The same model structure was applied for confidence judgments, with detection response included as an additional fixed-effects variable.

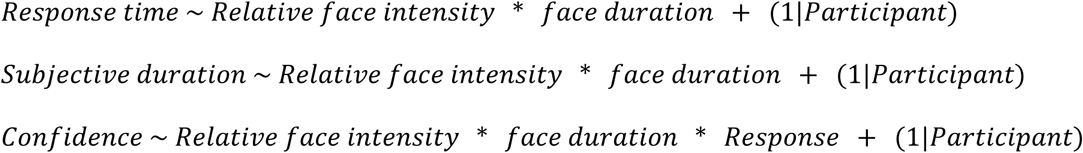

In experiment 3, we used the same regression models as in experiment 2, but we added the block type (predominantly short or predominantly long) as an independent variable in addition to face intensity and face duration.

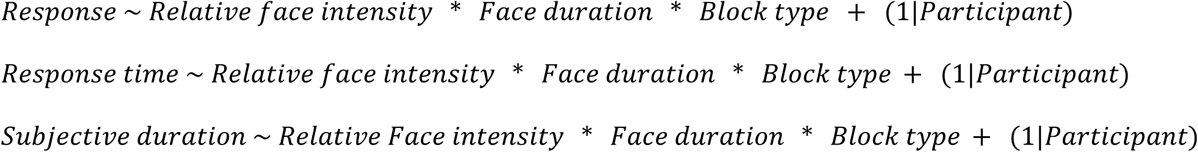

We added a random effect term for each model, including a by-participant (for experiments 2-3) or by-participant and session (for experiment 1) random intercept. Additionally, random slopes were added based on the model selection using the Bayesian information criterion (BIC).

### Computational modeling

We devised three models with different evidence accumulation mechanisms: One leaky evidence accumulation process (LEAP) with leakage as a free parameter, one perfect integrator with leakage fixed to zero, and an extrema model with an infinite leakage. Therefore, the perfect integrator integrates evidence perfectly, and the extrema model samples evidence sequentially from a stationary distribution without any integration (Stine et al., 2020). Models were compared based on the model evidence (evidence lower bound ELBO).

#### Leaky evidence accumulation process

The model involved a dynamic drift rate µ(*t*) defined as the mean of the noisy evidence being accumulated over time, a leakage parameter λ (free parameter) driving accumulated evidence towards zero, and additive white noise *W*(*t*) (**Eq. 1**). For the sake of biological plausibility, the evidence accumulation process was constrained to be positive, as firing rates are always positive. The accumulator was initialized at zero, and evolved according to:

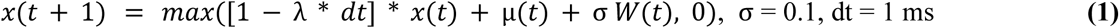

On stimulus onset, the accumulation process began accumulating the noise alone (µ(t)=0), until face onset plus a non-decision time *t_ndt_* (free parameter) *t*_*on*_ + *t*_*ndt*_, at which point µ(*t*) increased to a level Ɣ * *log*(*I* + 1), where *I* represents the stimulus intensity and Ɣ a scaling factor (free parameter). We used the logarithmic function to avoid −∞ values, as *I* included values of 0, 0.8, 1, 1.2, and 1.4. µ(t) then returned to zero at the same delay beyond face offset (*t*_*off*_ + *t*_*nd*_), thereafter continuing to accumulate only noise (**Eq.2**). Thus:

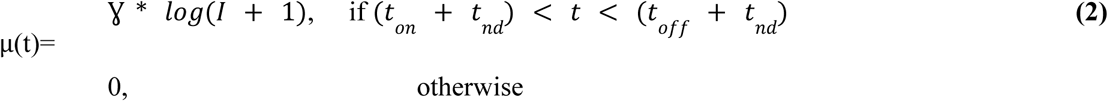

The event of committing to a face detection decision and its timing were determined by the accumulated evidence reaching a decision threshold *θ* (free parameter).

#### The perfect integrator

The perfect integrator consisted of a drift-diffusion model according to which samples of momentary evidence are perfectly accumulated over time, without any leakage. As for LEAP, on face onset, the drift rate rose to a level Ɣ * *log*(*I* + 1) for a duration *d* corresponding to the face’s physical duration. If the accumulated evidence (**Eq. 3**) reached the decision threshold, the face was considered as detected, otherwise, it was missed.

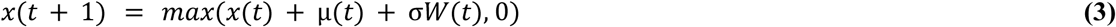

#### Extrema model

The extrema model involved no integration of evidence, i.e., each sample of evidence was independent of the other (**Eq. 4**). At each time step, the sample of momentary evidence *x*(*t*) was compared to the decision threshold *θ*. If the sampled evidence exceeded the threshold, the face was detected and the process was terminated otherwise, the sampled evidence was discarded. Evidence was sampled from the same signal as in LEAP and the perfect integrator model.

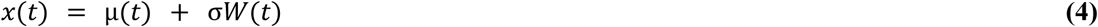

#### Confidence and subjective duration models

Subjective confidence *S*_*Conf*_ was evaluated by tracking the maximum distance δ between *x*(*t*) and the decision threshold over the trial and dividing it by the decision threshold value. We reasoned that this normalized readout δ would allow us to better compare model parameters across participants as the confidence readout is expressed in decision threshold units. We multiplied the resulting readout δ with a scaling factor α (confidence sensitivity) and a shifting factor β_*C*_ (confidence bias) to account for the fact that participants could have different confidence sensitivities and different over- or under-confidence biases (Fleming & Lau, 2014). The resulting value was then restricted to the 0-1 interval using a sigmoidal function (**Eq. 5**)

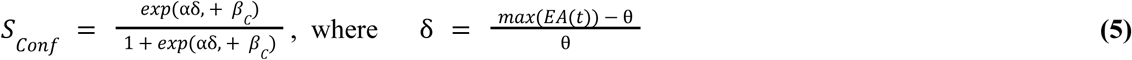

Subjective duration *S*_*Dur*_ was assessed as the time difference *t*_*diff*_ between the moment when *x*(*t*) first reached a perceptual threshold ɸ = **δ**θ (free parameter **δ** times the decision threshold θ), assumed to be lower than the decision threshold (**δ** <1), and the moment when it dropped below it. We also added a duration bias β_*D*_ to account for systematic tendencies to over- or under-estimate duration (**Eq. 6**).

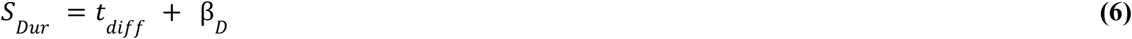

Because of stochasticity built into the evidence accumulation process, some trials had small periods of time during which accumulated evidence fell below the perceptual threshold and then rose above it again. We considered that the face would still be perceived during these gaps as long as they were inferior to 200 ms. There were only 1.29±176% of trials with longer gaps.

#### Fitting procedure

All three models were fitted to detection rates and response times across the 13 experimental conditions (a total of 3584 trials) using the Variational Bayesian Monte Carlo (VBMC) toolbox (Acerbi, 2020). This method approximates the posterior distribution of the model parameters and estimates the log model evidence (or the evidence lower bound ELBO). We computed the log-likelihood based on the *G*^2^ statistic.

For each condition, responses were categorized into eight groups: proportion of misses (trials where participants did not detect the face), proportion of anticipatory responses (trials with response times < 0), and proportions of positive response times divided into six bins defined by the 0%, 10%, 30%, 50%, 70%, 90%, and 100% quantiles of response time distribution. The *G*^2^ statistic was calculated as:

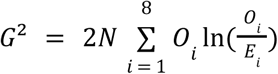

Here, N is the number of trials in each condition, *O*_*i*_ and *E*_*i*_ are the observed and simulated proportions, respectively, for a group *i*. The log-likelihood *L* of the model given the data is related to the *G*^2^ statistic by:

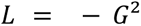

For confidence and subjective duration data, the log-likelihood was estimated by comparing the observed and simulated confidence and duration distributions using also the G-squared statistic. Confidence and subjective duration data for each experimental condition were divided into six bins corresponding to the 0%, 10%, 30%, 50%, 70%, 90%, and 100% quantiles of the observed and simulated distributions. Then, relative frequencies were calculated for observed and simulated data and the G-squared statistic quantifies the divergence between them, with the log-likelihood defined as the negative G-squared value. For each model, the fitting procedure was repeated five times with different starting points sampled from the prior and we selected the run providing the best ELBO.

#### Deviation from pre-registered plan

We followed the pre-registered plan except for not modeling the decay of sensory evidence over time using two parameters, as this led to non-identifiable models. Our final decisional model comprises four parameters instead of the six we had planned.

#### Parameter recovery

We performed a parameter recovery analysis by simulating 20 synthetic datasets (detection rates and response times distributions) using different values of decision threshold, drift rate, and leakage. We then fitted the LEAP model to each simulated dataset and calculated the correlation between the generator parameters and the recovered ones. We found a strong correlation between the generated and the recovered parameters (R ≥ 0.89; **Figure S14**).

#### Model recovery

We also performed a model recovery analysis to ensure that the three models used to fit detection rates and response times were distinguishable. To this end, we generated five synthetic datasets from each model using five different sets of parameters. Each data set was fitted with all models, and for each pair of simulated and fitting models, we calculated the proportion of times each model provided the best fit (based on the best ELBO). The results presented in the confusion matrix (**Figure S15**) indicate a successful recovery since the confusion matrix is predominantly diagonal. This diagonal pattern means that a given dataset was best fitted by its true generating model.

## Acknowledgments

The authors would like to thank Artemio Soto Breceda for the useful discussions and his valuable advice. NF has received funding from the European Research Council (ERC) under the European Union’s Horizon 2020 research and innovation program (Grant Agreement No. 803122). MP was funded/co-funded by the European Union (ERC, LEAP, 101077874). Views and opinions expressed are however those of the author(s) only and do not necessarily reflect those of the European Union or the European Research Council. Neither the European Union nor the granting authority can be held responsible for them.

## Authors’ contributions

Conceptualization and methodology: all authors; data collection: RM, ES; software and code validation: RM, MP; supervision: SK, NF, MP; funding acquisition: NF; original draft: RM, NF, MP; writing - reviewing and editing: all authors.

## Data availability statement

Data and code will be made public upon publication.

