## Supplementary information for "Leaky evidence accumulation accounts for perceptual confidence and subjective duration"

### Supplemental Information

#### S1. Experiment 2 results

In experiment 2, we tested our behavioral predictions on a larger cohort of healthy participants (N = 20). Participants' average detection rate was 67.66 % (standard deviation (s.d.) = 5.23). The false alarm rate was 8.94 % (s.d. = 8.34). As expected, detection rates were higher for static stimuli with higher intensities (β = 0.83; $p < 0.001$), but we found no effect of stimulus duration on detection (β = -3.85e-5; $p = 0.99$; Figure 2A) nor an interaction between intensity and duration (β = 0.08; $p = 0.14$). In the dynamic condition, stimuli with ramping-down intensity profile were better detected than stimuli with ramping-up intensity (β =-0.12; $p = 0.025$). However, there was no effect of stimulus duration (β = -0.08; $p = 0.1$) on detection rates. In line with our hypotheses, higher stimulus intensity (β = -0.05; $p < 0.001$) and shorter stimulus duration (β = 0.14; $p < 0.001$) led to faster response times for static stimuli (Figure 2B). For dynamic stimuli, ramping down intensities (β = 0.14; $p < 0.001$) and shorter stimulus durations (β = -0.07; $p < 0.001$) led to shorter response times, consistent with our pre-registered hypotheses.

Participants from experiment 2 were more confident as the stimulus intensity increased in the static condition (β = 0.04; $p < 0.001$), and this effect was significantly more pronounced for "yes" responses (when participants reported seeing a face) compared to "no" responses (when they did not respond) (static intensity x response interaction: β = -0.03; p < 0.01). When participants did not report perceiving a face, we observed no significant effect of stimulus intensity on confidence (β = 0.001; $p = 0.85$). Our analysis showed no effect of stimulus physical duration on confidence judgments, neither for yes (β = 0.01; $p$=0.22) nor for no responses (β = -0.03; $p = 0.07$). For the dynamic condition, we found no effect of the stimulus ramping profile (β = 0.0008; $p = 0.9$), stimulus duration (β = -0.01; $p = 0.18$), or participant responses (β = -0.04; $p = 0.05$) on confidence ratings (**Figure 3A**). These results for static stimuli aligned with our predictions based on a greater accumulation of evidence beyond the detection boundary for stimuli with higher intensity, whereas the results for dynamic stimuli deviated from our predictions, suggesting no effect of the intensity ramping profile on the maximum of accumulated evidence.

Consistent with our predictions, participants in experiment 2 reported longer durations for stimuli of higher static intensity, and this effect was larger for long stimuli (β = 0.15; $p < 0.001$) than for short ones (β = 0.08; $p < 0.001$), with no significant interaction effect observed (β = 0.008; $p = 0.5$). In the dynamic condition, the subjective duration was affected by the intensity variation profile (β = -0.04; $p = 0.016$) and the actual stimulus duration (β = -0.1; $p < 0.001$), indicating that long stimuli and stimuli with ramping-down intensities were perceived as lasting longer. These findings supported our predictions that a higher drift rate results in the accumulated evidence remaining above the decision threshold for a longer time, implying a longer subjective duration.

#### S2. Speed accuracy tradeoff of the leakage parameter

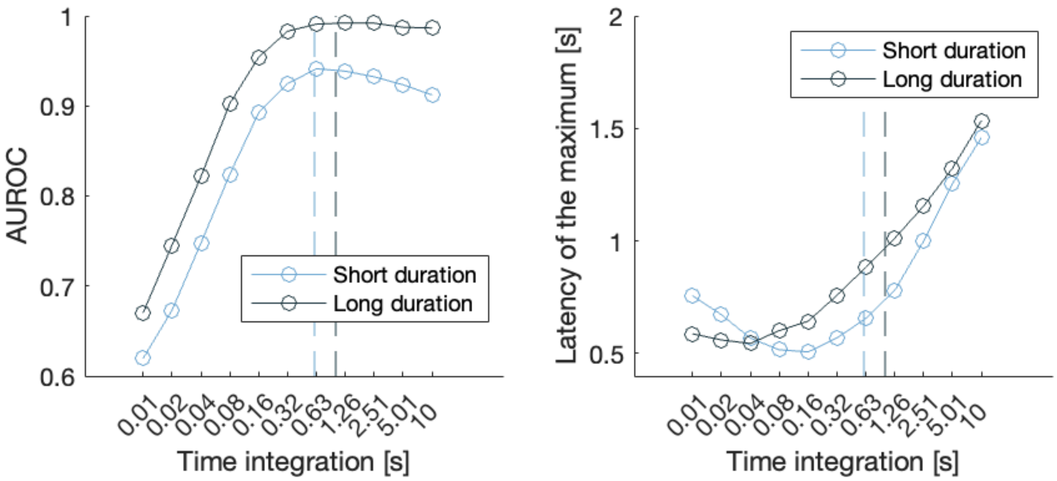


*Figure: Model simulations showing the tradeoff between reactiveness and optimal discrimination between noise (catch trials) and signal trials. Leakage values were varied in order to obtain integration windows ranging from 10 ms to 10 s. Decision threshold was set to 1.5, drift rate to 0.02, and non-decision time was zero.* ***A.*** *The Area Under the Receiver Operating Characteristic curve (AUROC), which quantifies the model's ability to discriminate between noise (catch trials) and signal presented at the liminal intensity as a function of the integration window for short (light blue curve) and long (deep blue curve) stimuli. The two vertical dashed lines indicate the optimal integration windows for detecting short (light blue) and long (deep blue) stimuli, representing the optimal point where too little leakage leads to excessive false alarms and too much leakage results in too many misses. Note that with higher drift values, the AUROC curve plateaus.* ***B.*** *Timing of the maximum accumulated evidence as a function of the integration window for short (light blue curve) and long (deep blue curve) stimuli. The two vertical dashed lines indicate the same optimal integration windows for short (light blue) and long (deep blue) stimuli as panel A, showing that shorter windows provide a relative benefit in timing.*

#### S3. Evidence accumulation traces

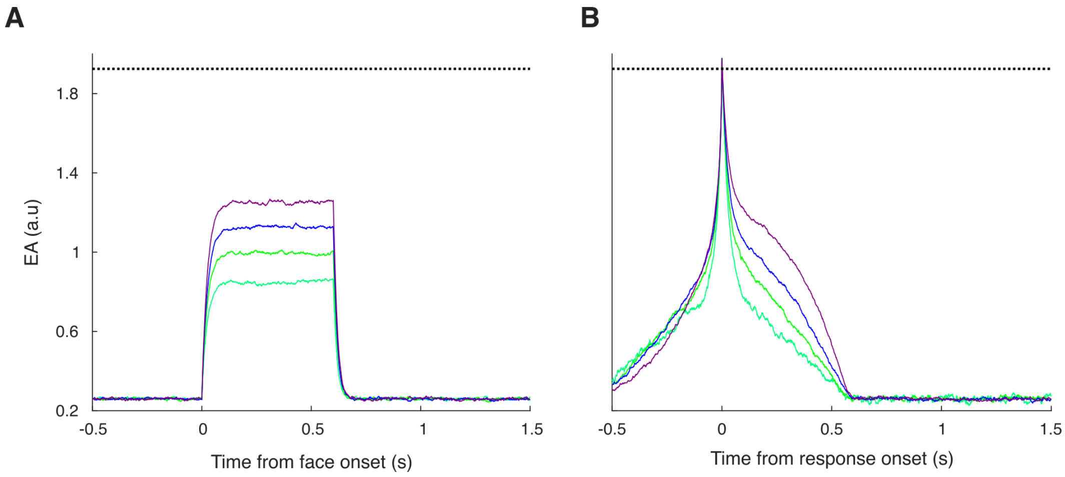


*Figure: Traces of evidence accumulation for one participant, averaged across the four static intensity conditions. The dotted line represents the detection threshold. Traces for the other three participants exhibit similar patterns to those shown here.* ***A****. Traces time-locked to face onset. Note that while the average accumulated evidence remains*

*under the decision threshold, this does not prevent individual traces from reaching it at varying latencies.* ***B****. Traces time-locked to response onset showing how higher intensities lead to longer perceived duration.*

### S4. Perfect integrator fitting results for detection

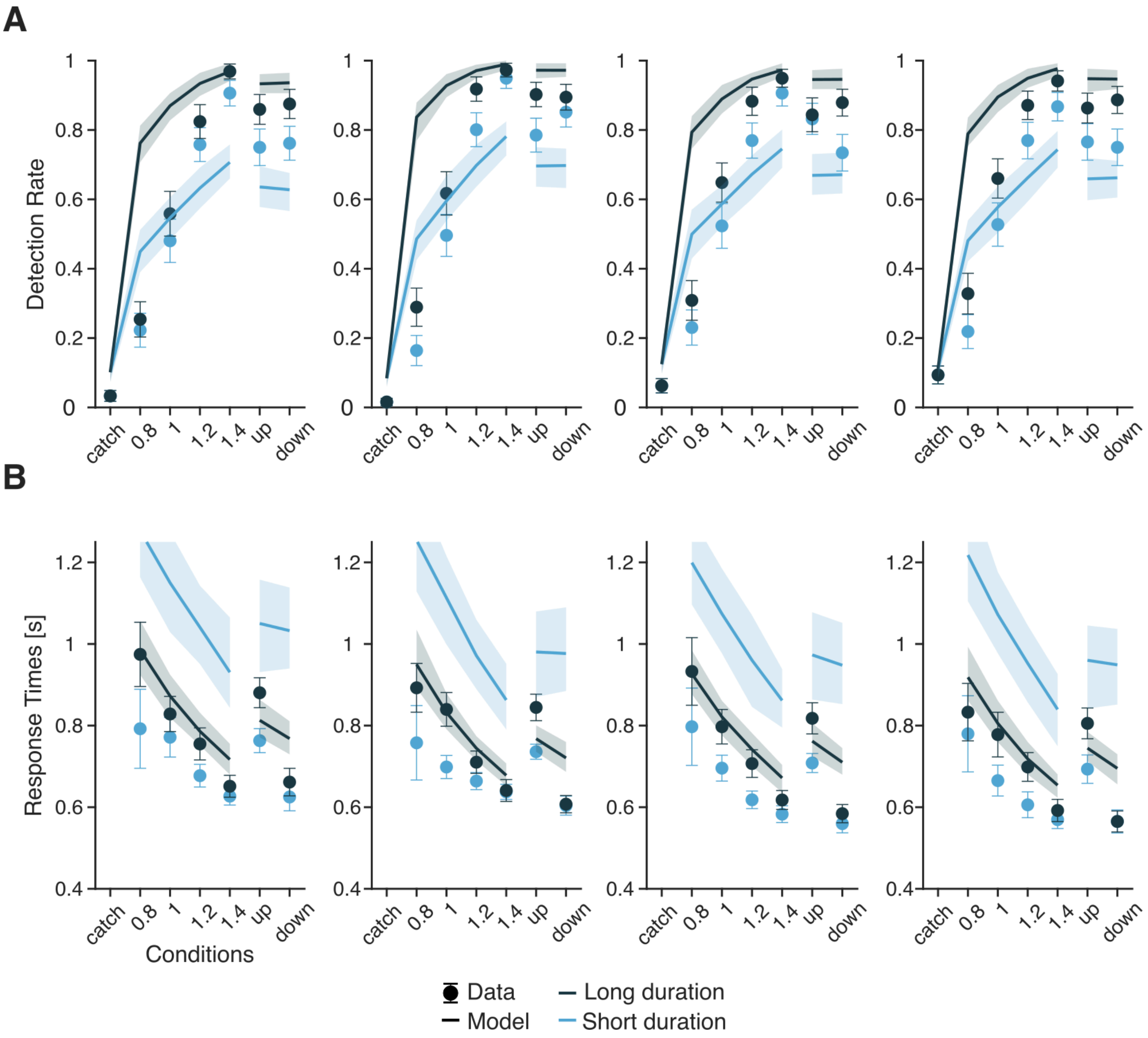


*Figure: Detection results in experiments 1 and 2 with perfect integrator model fits.* ***A****. Individual detection rates for each of the four participants in experiment 1 (light and deep blue dots for short and long faces, respectively). Solid lines represent model fits for short faces (light blue traces) and long faces (deep blue traces), with shaded areas indicating 95% confidence intervals.* ***B****. Average detection rates across 20 participants for long faces (deep blue squares) and short faces (light blue squares) as a function of stimulus conditions in experiment 2.* ***C****. Individual response times in experiment 1. Solid lines represent model fits for short faces (light blue traces) and long faces (deep blue traces), with shaded areas indicating 95% confidence intervals.* ***D****. Averaged response times in experiment 2 for long faces (deep blue line) and short faces (light blue line) as a function of stimulus conditions.*

**S5. Extrema model fitting results for detection**

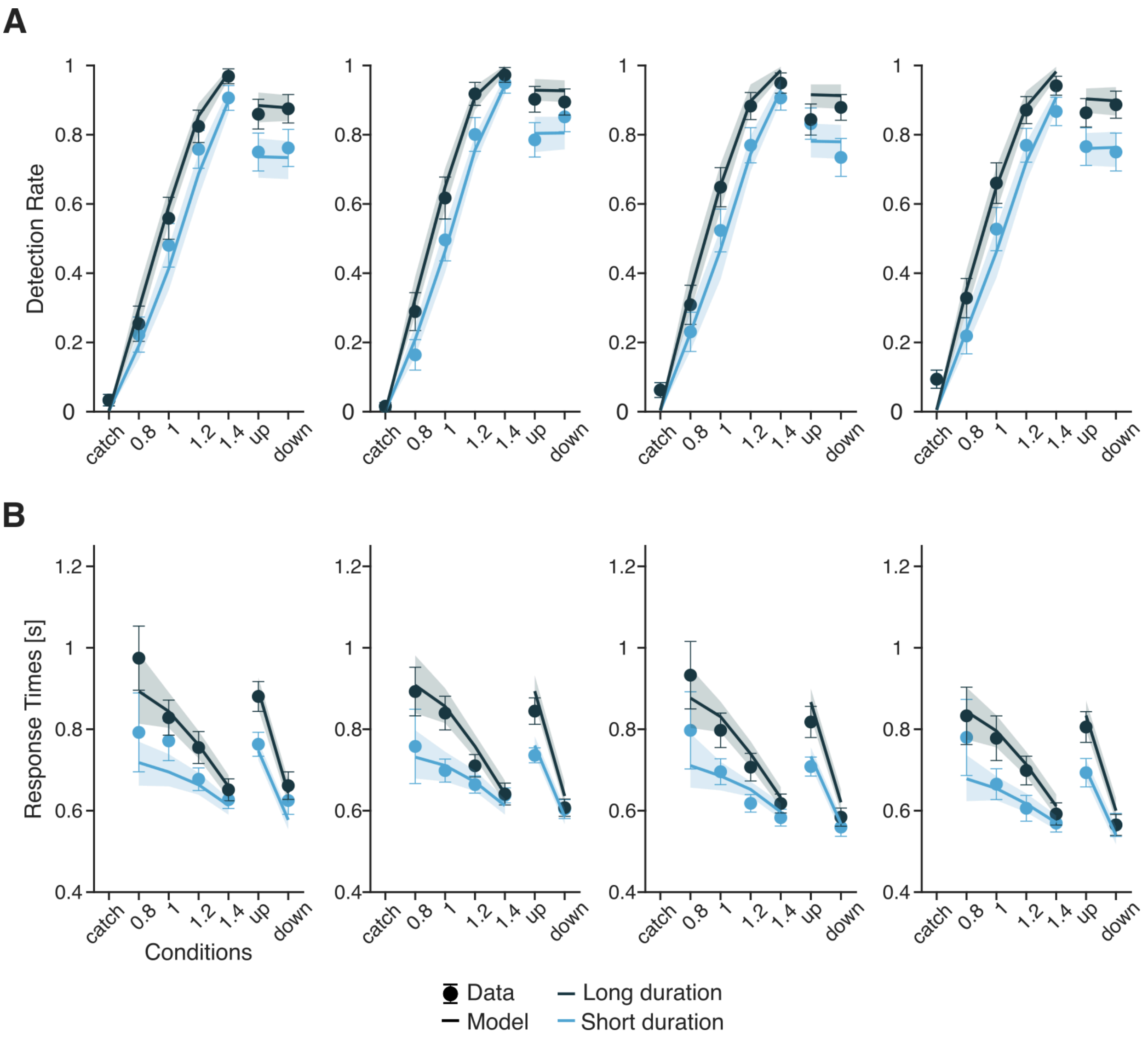


*Figure: Detection results in experiments 1 and 2 with extrema model fits.* ***A****. Individual detection rates for each of the four participants in experiment 1 (light and deep blue dots for short and long faces, respectively). Solid lines represent model fits for short faces (light blue traces) and long faces (deep blue traces), with shaded areas indicating 95% confidence intervals.* ***B****. Average detection rates across 20 participants for long faces (deep blue squares) and short faces (light blue squares) as a function of stimulus conditions in experiment 2.* ***C****. Individual response times in experiment 1. Solid lines represent model fits for short faces (light blue traces) and long faces (deep blue traces), with shaded areas indicating 95% confidence intervals.* ***D****. Averaged response times in experiment 2 for long faces (deep blue line) and short faces (light blue line) as a function of stimulus conditions.*

#### S6. Models comparison results

| Participants | Perfect Integrator | Extrema model | LEAP |
|---|---|---|---|
| P1 | -6106 | -859.33 | -867.55 |
| P2 | -7196.8 | -593.21 | -461.73 |
| P3 | -5985.4 | -623.94 | -538.19 |
| P4 | -5188.7 | -255.39 | -264.12 |

*Table: Values correspond to the Evidence Lower Bound (ELBO). Higher ELBO values are considered as support for the best model.*

#### S7. Best fitting detection parameters for each participant

| Participants | Decision threshold | Drift rate | Leakage ($s^{-1}$) | Non-decision time (s) |
|---|---|---|---|---|
| P1 | 1.93 | 0.0536 | 37.39 | 0.383 |
| P2 | 2.79 | 0.0501 | 20.23 | 0.373 |
| P3 | 1.74 | 0.0565 | 44.15 | 0.378 |
| P4 | 1.81 | 0.0499 | 38.02 | 0.336 |

*Table: Individual fitted parameters for the leaky evidence accumulation model. Values correspond to the mode of the posterior distribution for each parameter.*

### S8. Confidence in “no” responses in experiments 1 and 2

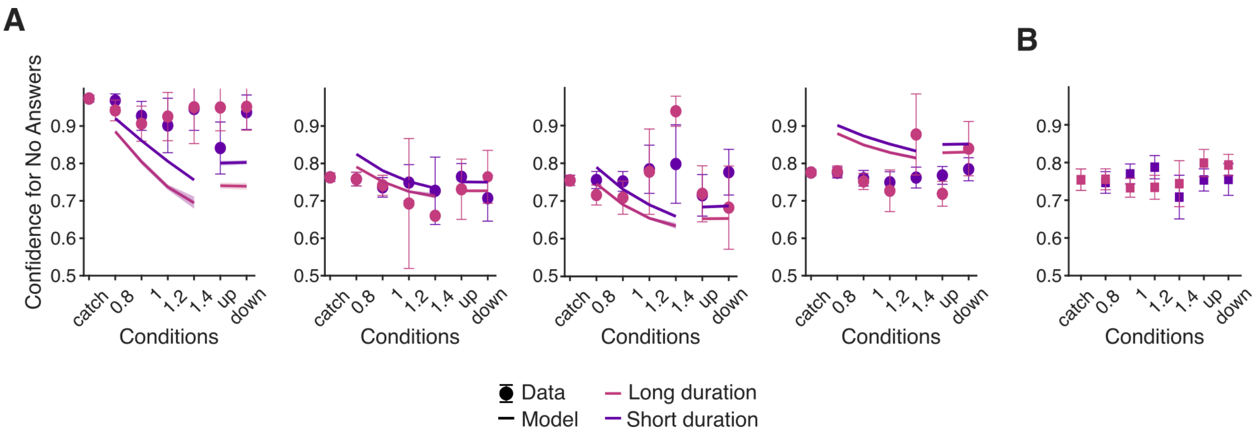


*Figure: Confidence for “no” answers in experiments 1 and 2. **A**. Individual confidence judgments for “no” answers from four participants in experiment 1 (deep purple and pink dots for short and long stimuli, respectively). Solid lines represent model fits for short stimuli (deep purple traces) and long stimuli (pink traces), with shaded areas indicating 95% confidence intervals. **B**. Average confidence judgments for “no” answers across 20 participants for long stimuli (pink squares) and short stimuli (deep purple squares) as a function of stimulus conditions in experiment 2.*

### S9. Best fitting confidence parameters for each participant

| Participants | Confidence sensitivity | Confidence bias |
|---|---|---|
| P1 | 28.28 | 0.12 |
| P2 | 6.73 | 0.74 |
| P3 | 10 | 0.31 |
| P4 | 9.63 | 1.25 |

*Table: Individual fitted parameters for confidence ratings. Values correspond to the mode of the posterior distribution for each parameter.*

### S10. Distributions of confidence ratings and model fits

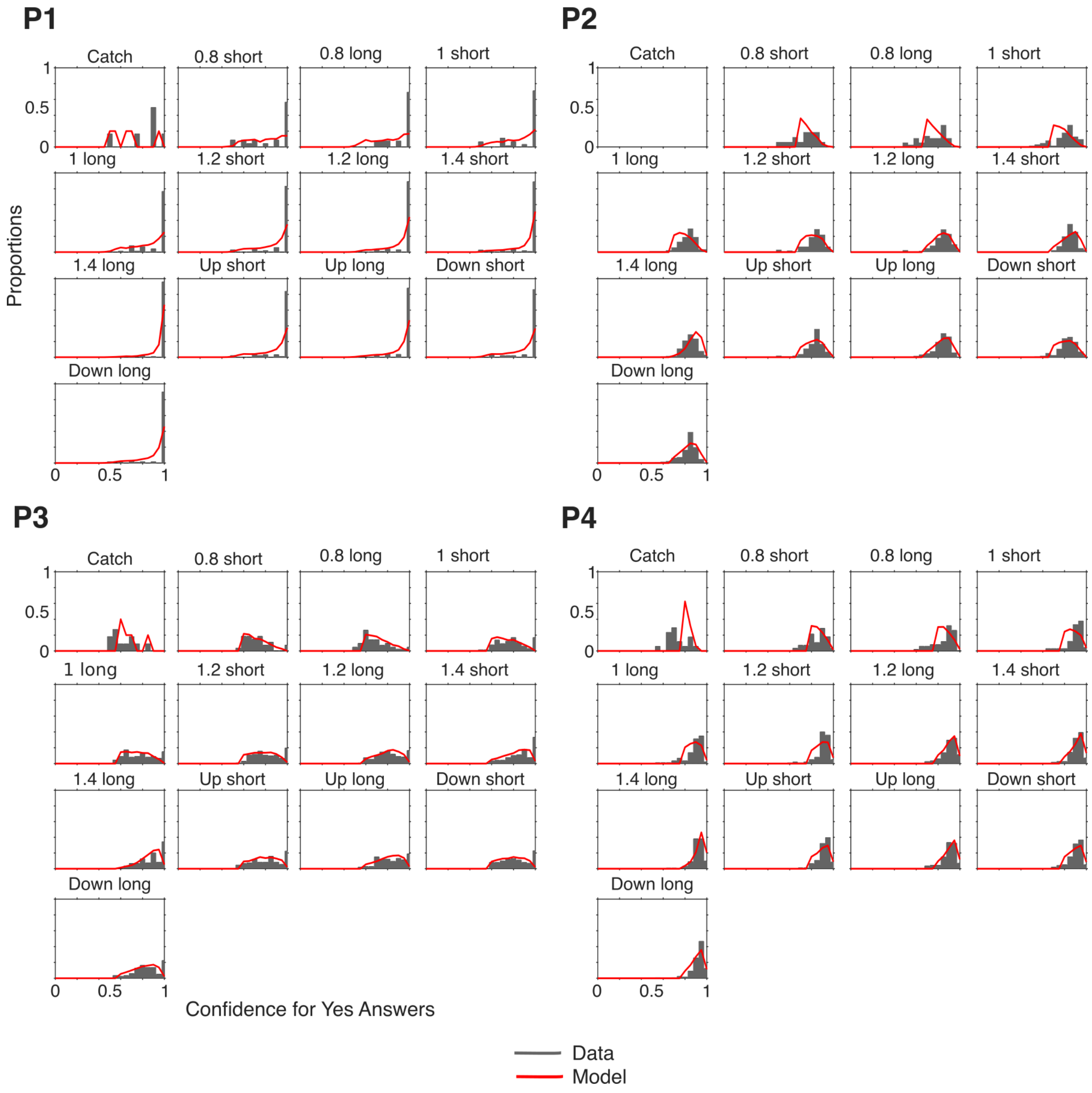


*Figure: Distributions of confidence ratings for detected stimuli in experiment 1, shown for four participants across each experimental condition and stimulus duration. Dark gray histograms represent behavioral data, while red lines indicate model fits.*

### S11. Effect of response times on confidence and perceived duration

We conducted a regression analysis to examine the effect of response times on confidence judgments. We found that in both static and dynamic conditions, faster responses were associated with higher confidence judgments ($\beta = -0.16$; $p < 0.001$ for the static condition, and $\beta = -0.17$; $p < 0.001$ for the dynamic condition).

Another regression model showed a significant effect of response times on perceived duration, suggesting longer subjective duration for shorter response times. We found this effect in both static ($\beta = -0.38$; $p < 0.001$) and dynamic ($\beta = -0.52$; $p < 0.001$) conditions.

The effect of response times on confidence judgments and subjective duration was also significant in model simulations for all participants ($p < 0.001$).

### S12. Best fitting duration parameters for each participant

| Participants | Perceptual threshold | Duration bias |
|---|---|---|
| P1 | 0.84 | 0.15 |
| P2 | 1 | 0.3 |
| P3 | 0.85 | 0.44 |
| P4 | 0.83 | 0.5 |

*Table: Individual fitted parameters for the subjective duration. Values correspond to the mode of the posterior distribution for each parameter.*

### S13. Distributions of subjective duration and model fits

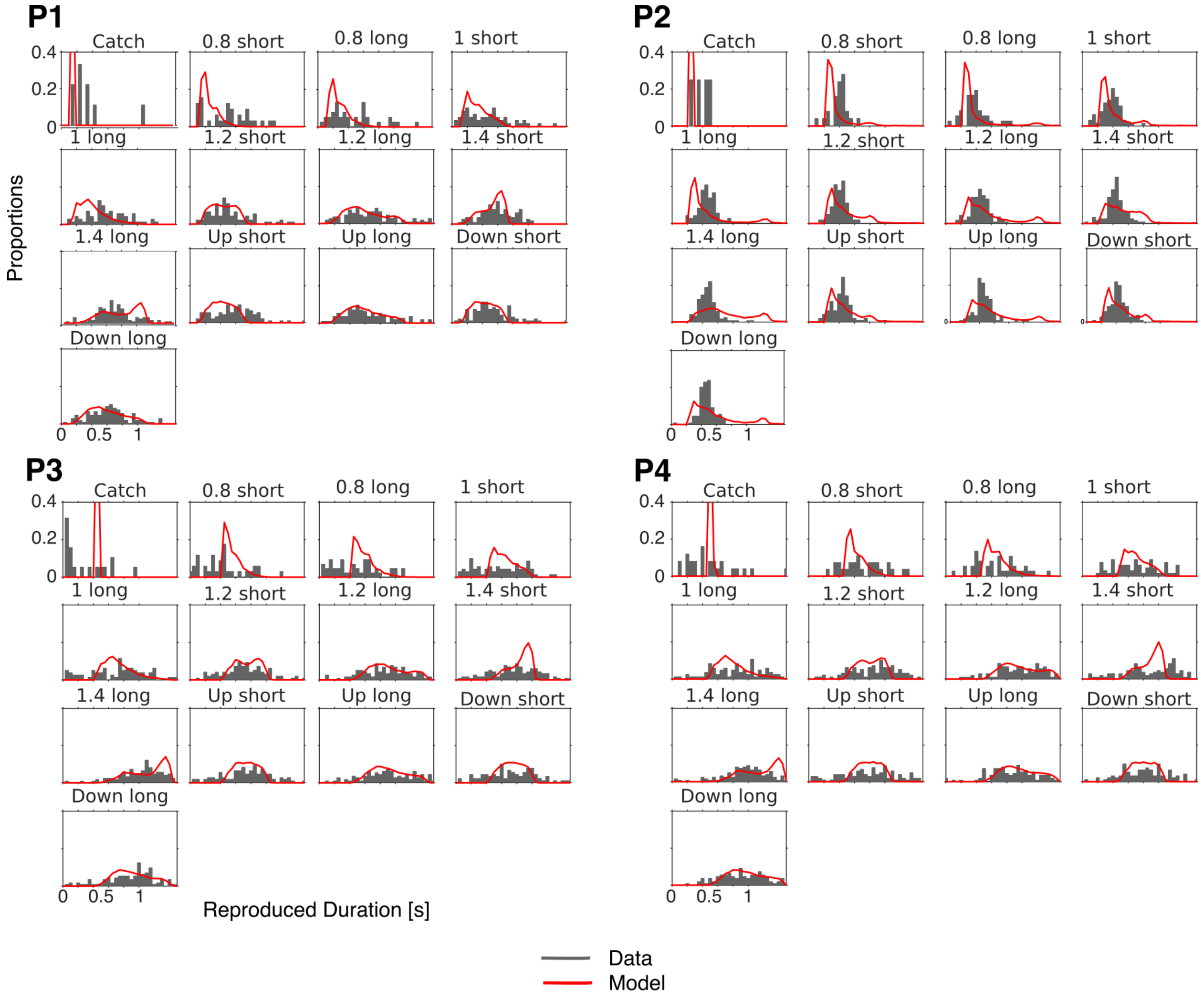


*Figure: Distributions of reproduced durations in experiment 1, shown for four participants across each experimental condition and stimulus duration. Dark gray histograms represent behavioral data, while red lines indicate model fits.*

### S14. Parameters recovery results

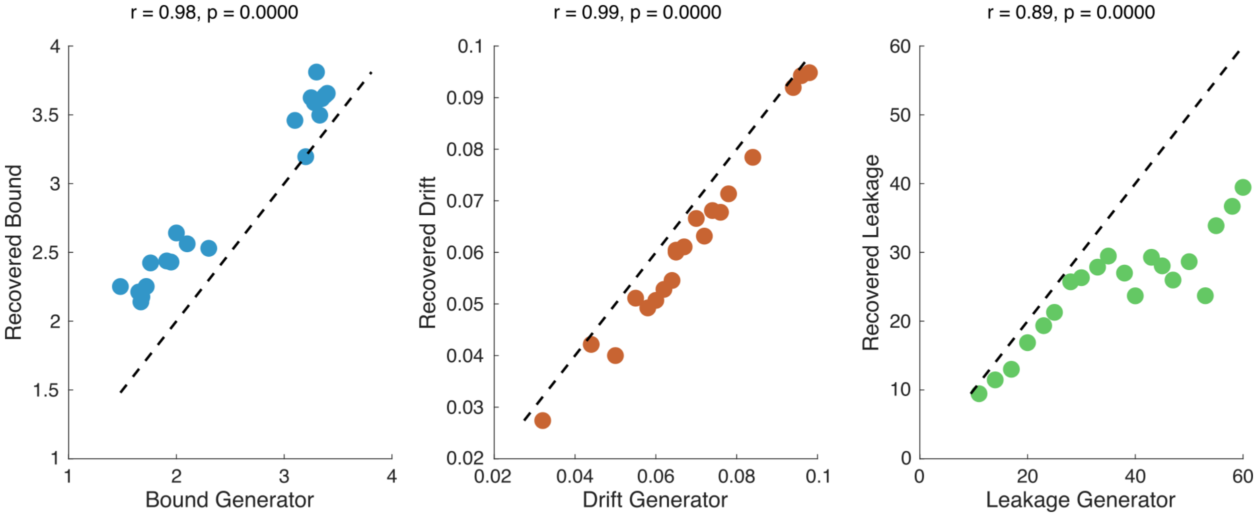


*Figure: Correlation between the generated and recovered parameters for the decision threshold, the drift rate, and the leakage. Each subplot shows a scatter plot of generator versus recovered values (blue dots), with a red line representing a linear fit to the data points. Correlation coefficients (r) and p-values (p) are included to assess the accuracy of the recovery process, demonstrating a strong correlation between the generated and recovered values for all three parameters.*

### S15. Model recovery results

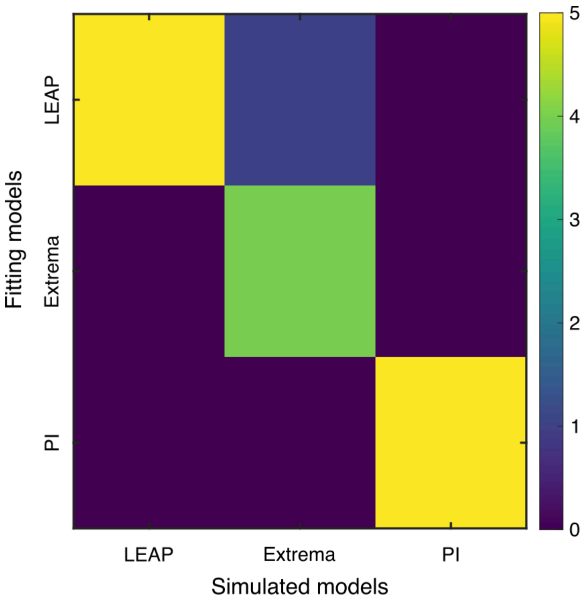


*Figure: Five synthetic datasets were generated from each model using different parameter sets. Each dataset was fitted with all three models, and the proportion of times each model outperformed the others (based on the best ELBO) was computed. The color intensity shown in this matrix indicates the proportion of "wins" for each fitting model. The predominantly diagonal pattern demonstrates successful recovery.*
